# A dynamic repertoire of wound closure strategies precedes whole-body regeneration

**DOI:** 10.1101/2025.02.03.636261

**Authors:** Allison P. Kann, Mansi Srivastava

## Abstract

Wound closure is an essential aspect of successful regeneration, often acting as the first morphogenetic event that precedes downstream cellular events. Despite heavy investment in studying whole-body regeneration in many invertebrate systems, the steps by which these organisms heal their wounds remain understudied. Here, we investigate the cellular mechanisms of wound healing in the acoel *Hofstenia miamia*, an invertebrate worm capable of whole-body regeneration. *H. miamia* have two distinct epithelial layers, an outer epidermis and the epithelial lining of their pharynx. By labeling regenerating fragments with an actin dye, we found that *H. miamia* use distinct mechanisms of epithelial wound repair across different injury contexts. In transverse wounds that don’t injure the pharyngeal epithelium, the epidermis closes by gradual radial constriction. In contrast, injuries that damage both the epidermis and the pharyngeal epithelium show the formation of long, actin-rich protrusions that cross the wound gap and form heterotypic bridges prior to re-epithelialization. Muscle contraction is required for the formation of heterotypic bridges - when animals are anesthetized and immobile, they are unable to form these cellular bridges, and epidermal cells cannot migrate independently. Global actomyosin contractility also plays a role in repair mechanisms, and pharmacological perturbation of contractility shifts the dynamics of wound closure after amputation. In the presence of blebbistatin, heterotypic bridge formation is inhibited but homotypic re-epithelialization is accelerated through increased cell crawling. Together, this work identifies mechanisms by which epithelial layers close large wounds *in vivo*, identifying novel heterotypic cellular bridges in wound closure and demonstrating the conservation of actin-mediated processes in an early-diverging phylum.

## INTRODUCTION

The ability of animals to regenerate varies substantially across phyla, ranging from more limited, tissue-specific or structural regeneration to extreme cases of whole-body regeneration (WBR), where any missing cell or tissue type can be replaced after amputation^1,2^. Highly regenerative animals undergoing WBR face unique healing challenges and must be able to scarlessly repair virtually any injury prior to downstream regeneration. Most mechanistic studies of wound closure have focused on relatively simple injuries (i.e. damaging only the epidermis) in animals that are not capable of substantial regeneration, generating foundational knowledge about cellular behaviors but lacking a strong connection to downstream regeneration^3–8^. The work on wound closure in regenerative animals has primarily focused on either the formation and function of the ‘wound epidermis’, a specialized epidermis that covers the injury site and acts as an organizing center to induce downstream regeneration^9–12^, or large-scale transcriptional atlases detailing the molecular wound response^13–18^. Some studies have demonstrated the presence of conserved cellular mechanisms in regenerative animals^19–25^, but it remains ambiguous how much flexibility exists in closure strategies prior to regeneration. Specifically, it is unclear whether highly regenerative animals employ a uniform healing process regardless of injury context (i.e. major vs. minor wounds, varying geometries and tissue organizations) or if they have a range of mechanisms that can be tailored to different injury conditions.

Previous work has shown the presence of shared mechanisms of wound repair across the metazoan tree of life, including in early-branching lineages such as cnidarians^14,16,19,20^, and in protostomes^3,4,21,25–27^ and deuterostomes^15,18,28–30^. This suggests a deep conservation of cellular behaviors, evolving prior to the emergence of bilateral symmetry. A fundamental aspect of wound closure and subsequent healing is the process of re-epithelialization, in which a damaged epithelial layer undergoes cellular and morphological changes to re-establish an intact tissue^6^. Wound closure through cell crawling is facilitated through the formation of actin-based membrane protrusions (such as lamellipodia or filopodia) in cells at the periphery of a wound that drive migration into the gap^7,31,32^. Actomyosin contractility is essential in this crawling mechanism for both retraction at the rear of migrating cells and the formation of tension between adhesion points^19,33,34^. Purse string closure is similarly driven by contractility; cells around the wound form a supracellular cable, contracting and bringing the edges of cells inwards to pinch off the gap^31,35,36^. These mechanisms are not mutually exclusive and often work in tandem, bringing the edges of epithelial cells together across the open wound and restoring tissue integrity through the reestablishment of cell junctions^7^.

Studies both *in vivo* and *in vitro* have revealed that the mechanisms of re-epithelialization vary substantially between wounding conditions, with factors such as wound size, geometry, tissue type, extracellular matrix composition, and life stage of the animal all influencing the mechanism of closure^6,7, 19,23,26,41,33,38–41^. Large injuries typically involve damage to the underlying extracellular matrix, leading to either wound closure through the formation of actomyosin purse strings, which do not require an intact substrate, and/or the deposition of a transient matrix that provides a substrate for epithelial cell crawling^19,22,24,37,42^. These large injuries also require coordination between layers of cells, often relying on contraction of mesenchymal cells underneath the epithelium to drive closure^29,43,44^. Despite evidence of variable cellular behaviors across wound conditions, most *in vivo* studies have focused on factors that determine whether wounds close or not, rather than the flexibility of large-scale closure strategies within the same system. This is likely due to a few factors. First and foremost, traditional model organisms that have extensive genetic and imaging tools (e.g. *D. melanogaster, M. musculus*) have distinct physiological considerations. Not only are these species incapable of whole-body regeneration, restricting the types of injuries the animal can recover from, but they possess circulatory and immune systems that complicate and contribute to the healing process^44,45^. Second, visualizing wound closure in a quantifiable and reproducible manner typically requires live imaging, which is more feasible to set up for minor wounds (such as a scratch assay, as used in *C. hemisphaerica*^19^) than for large injuries. We therefore have a detailed understanding of some cellular decision-making, like switching between cell crawling and purse string formation depending on substrate^3,19,35,40,46^, but less is understood about whether animals utilize the same toolkit of closure strategies across extreme amputations. Studying these cellular processes in nontraditional animals offers a new approach to understanding fundamental aspects of biology, capitalizing on their regenerative properties to ask questions that are infeasible in other species.

*Hofstenia miamia* is a marine invertebrate worm belonging to the enigmatic and understudied phylum Xenacoelomorpha, which likely represents an early-diverging animal lineage that is a sister group to all other bilaterians (Fig. 1A-B)^47–49^. *H. miamia* has emerged as a new model system, largely studied for its ability to undergo whole-body regeneration. Like other animals capable of WBR, such as platyhelminthes, *H. miamia* rely on a population of adult pluripotent stem cells to drive regeneration ^50,51^. However, even in conditions that block regeneration (e.g. functional knockdown of required genes or irradiation of proliferative cells^47,50^), damaged fragments are still able to visibly close their wounds – suggesting that healing and regeneration are distinct processes. In addition to their remarkable regenerative ability, *H. miamia* possess a ‘tube within a tube’ anatomy, with an outer epidermis and a centrally located pharynx lined with an epithelium^52^. This dual-epithelial organization prompts both questions of how complex injuries can heal and how tissues maintain their identity during repair and regeneration.

**Figure 1.**
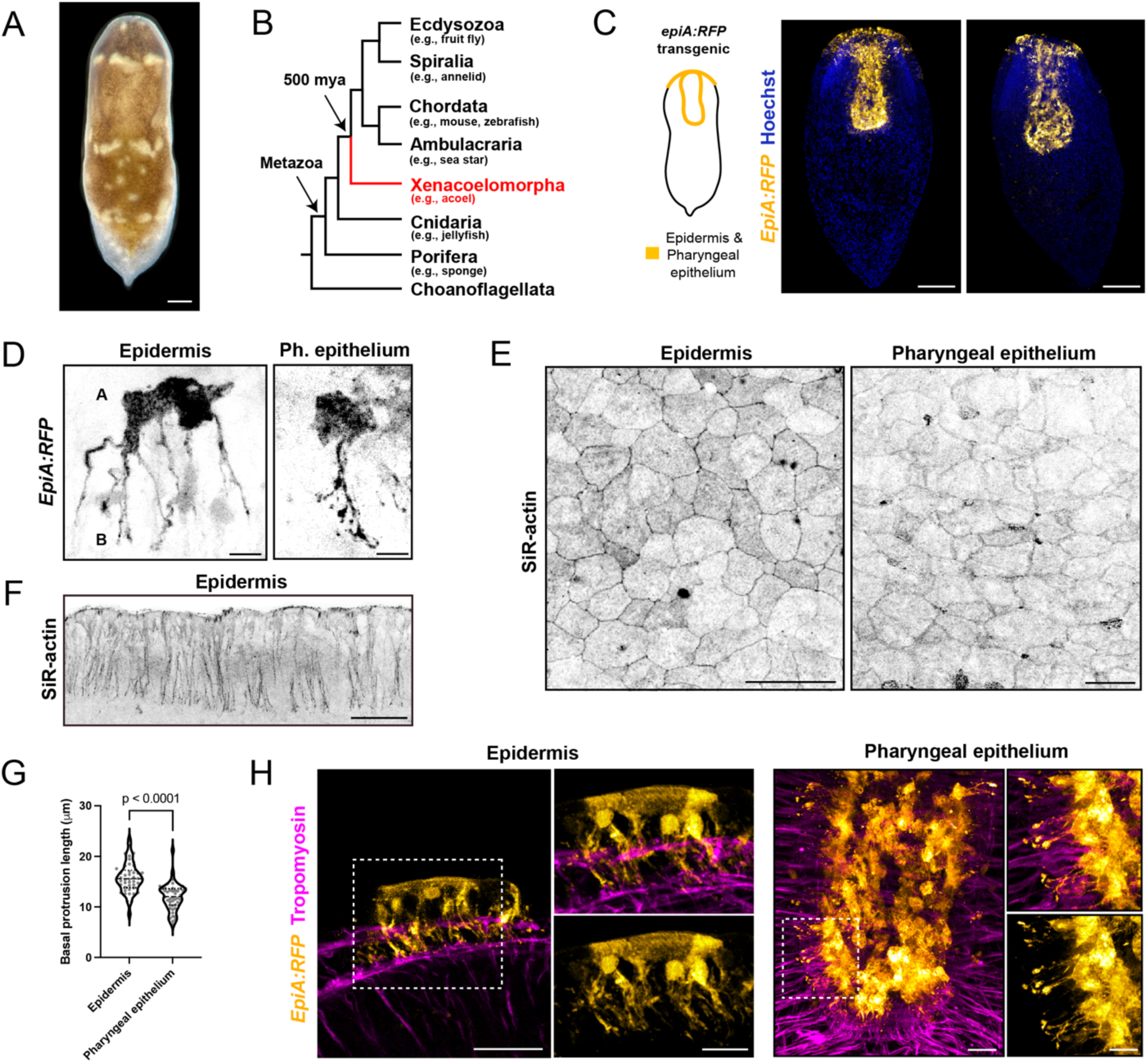
*Hofstenia miamia* have two distinct, atypical epithelial tissues. (A) A juvenile *Hofstenia miamia.* (B) A simplified phylogenetic tree showing the position of *Hofstenia miamia* (red, Xenoacoelomorpha) relative to other clades. mya = million years ago. (C) *EpiA:RFP* transgenic worms, fluorescently labeling the outer epidermis and the pharyngeal epithelium in a mosaic manner. (D) Morphology of individual epidermal and pharyngeal (Ph.) epithelial cells, visualized by *EpiA:RFP* transgenic labeling. A: apical, B: basal. (E-F) SiR-actin staining showing the apical organization of *H. miamia* epithelial tissues, particularly the polygonal shape of each cell and the presence of actin at cell-cell junctions (E) and the presence of actin within epidermal basal protrusions (F). (G) Quantification of basal protrusions within epidermal and pharyngeal epithelial cells. Data represent n=45 cells per cell type, pooled from 15 worms. Comparison by unpaired *t*-test. (H) Confocal images showing the proximity of muscle (visualized through immunofluorescence staining for tropomyosin) to both epithelial layers (visualized with EpiA:RFP transgenic labeling). Scale bars: 50 µm (A), 100 µm (C), 5µm (D), 20 µm (E, H), 25µm (F), 10 µm (H, zoomed-in box).

In this study, we investigate the healing capacity of *H. miamia*, examining whether wound repair mechanisms shift in response to different injury conditions. In relatively simple injuries (e.g. transverse amputations that only damage one epithelial layer, scratch wounds), animals rely on conserved mechanisms of epithelial contraction and migration. In more complex wounds (e.g. asymmetric sagittal wounds and other amputations that damage both epithelial layers), *H. miamia* form novel heterotypic bridges between epithelial layers prior to homotypic re-epithelialization. These bridges are dependent on both global actomyosin tension and contractility generated by neighboring muscle, but the regenerating fragments can modify their repair strategies in the face of mechanical perturbation. *H. miamia* therefore possess an adaptable toolkit of wound closure mechanisms, with flexible strategies to ensure successful repair and downstream regeneration regardless of injury type.

## RESULTS

### *Hofstenia miamia* have two distinct, atypical epithelial tissues

The acoel *Hofstenia miamia* can recover from virtually any injury throughout its lifespan, scarlessly healing wounds and rebuilding lost tissues^47,48^. Despite wound closure being an essential prerequisite to this extensive regeneration, the mechanisms of wound closure have not yet been studied in *H. miamia*. To fully understand the dynamics of re-epithelization, we first needed to characterize the epithelial layers of *H. miamia*.

*H. miamia* have two distinct epithelial tissues: (1) an outer epidermis that covers the whole animal, and (2) the inner lining of the pharynx, a tube-shaped organ that extends ∼1/2 down the anterior-posterior axis (Figure 1C). Although the dynamics of these cells have not yet been studied, previous transgenic labeling has revealed their cellular morphologies^52^. This transgenic labeling (using *EpiA:RFP* animals^52^) enables the visualization of detailed cell morphologies (Figure 1D), but the mosaicism prevents consistent imaging across animals. We found that the cytoskeletal dye SiR-actin robustly labels both layers of epithelial cells in fixed animals and can therefore be used to visualize entire tissues instead of individual cells (Figure 1E-F). The apical side of *H. miamia* epithelial cells resemble epithelia in other systems^53^, with the cells organized in a polygonal pattern on the surface of the animal (Figure 1E). As previously reported, the basal side of both epidermal and pharyngeal epithelial cells form cytoplasmic protrusions that extend into the body of the worm (Figure 1F-H). The protrusions are irregularly spaced and heterogeneous in number between cells but extend approximately 10-25µm in length, with the epidermal protrusions being slightly (but significantly) longer than those in the pharyngeal epithelium (Figure 1G). The basal protrusions in both cell types end in close proximity to underlying muscle (Figure 1H), but it is unclear whether cell-cell junctions form at the tips of these projections or whether they adhere to an atypical basement membrane.

The unique morphology and organization of these tissues raise questions of how epithelial cells mobilize to close wounds upon amputation and facilitate downstream regeneration. Even though *H. miamia* can recover from virtually any injury configuration, not all injuries require the same cellular response. Notably, the internal localization of the pharynx means that some amputations will damage both epithelial layers, adding another layer of complexity to the re-epithelialization process. We therefore sought to understand the dynamics of re-epithelialization during wound closure, comparing injuries that damage one or both epithelial layers.

### Amputations that damage the outer epidermal layer close through actin-based constriction

Transverse amputations through the middle of the animal, separating the head from tail, have been utilized as the dominant assay for whole-body regeneration across systems^14,47,48,50,51^. In *H. miamia,* a typical transverse amputation aims to cut at the middle pigmentation stripe observable on the dorsal surface of the worm, leaving behind a large gap with open epidermal edges around the wound. This amputation halves some tissues (e.g. gut, nerve net, and musculature) and completely removes anterior organs (e.g. pharynx and brain) from the posterior fragment. In addition to the biological process of wound healing, these fragments therefore face different regenerative challenges: the head fragment needs to regenerate the posterior tissues, whereas the tail fragment must regenerate anterior-specific tissues and organs (Figure 2A).

**Figure 2.**
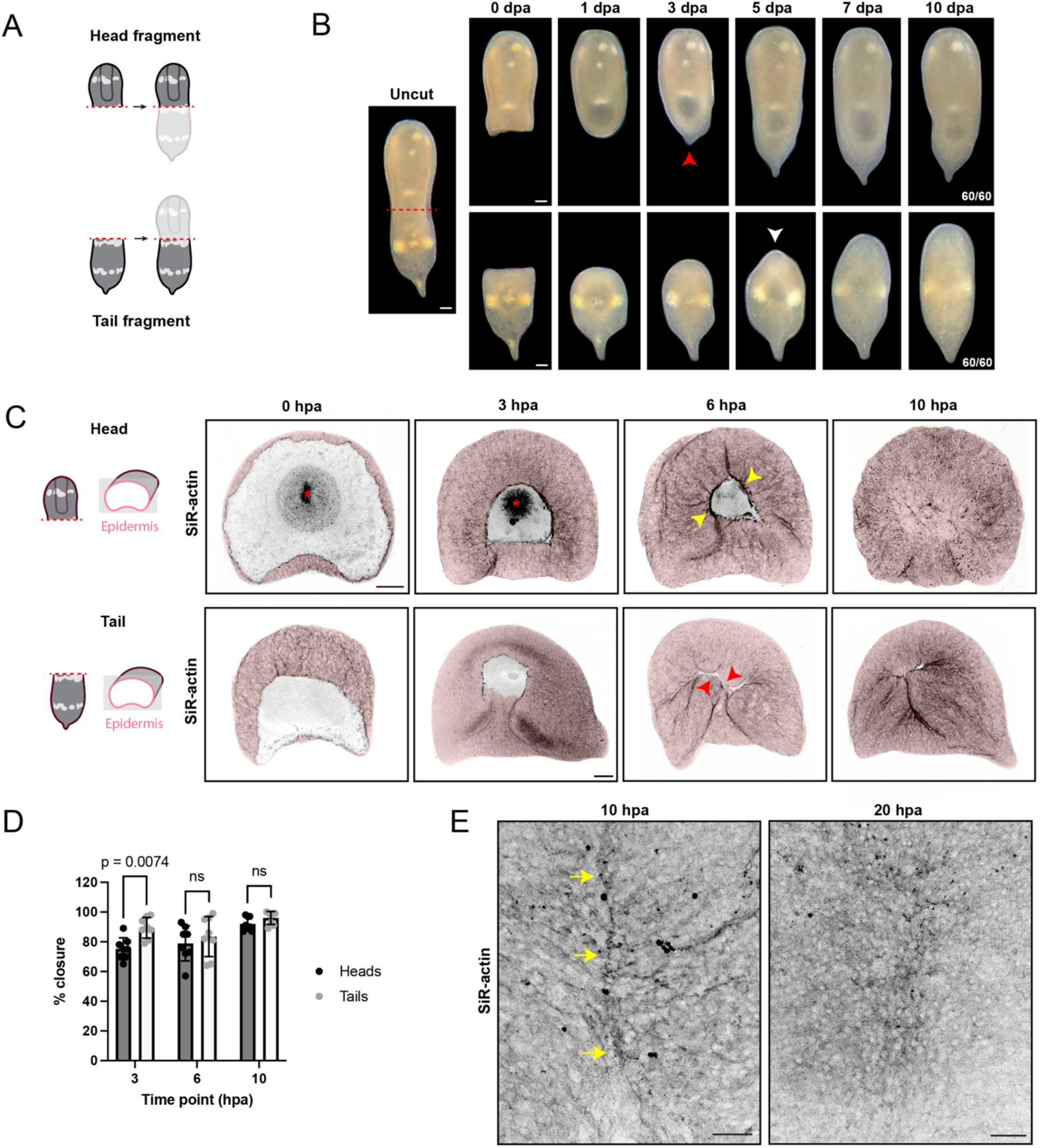
Transverse amputations close through gradual constriction of the epidermis across the wound edge. (A-B) Schematic (A) and representative brightfield time course (B) showing regeneration of *H. miamia* following transverse amputation. Red arrowhead indicates the formation of a pointed tail, white arrowhead indicates a visible blastema. (C) Representative images of wound closure following transverse amputation between 0-10 hours post-amputation. Red asterisk indicates the base of the pharynx, visible in open wounds. The outer epidermis is pseudocolored in pink for clarity. (D) Quantification of wound closure following transverse amputation, comparing between head and tail fragments from 3-10 hours post-amputation. Data represent 8 worms per time point, and comparisons were performed by 2-way ANOVA with Šidák’s multiple comparisons test; ns=not significant. (E) High-magnification images showing continued dynamics of re-epithelialization following initial closure; yellow arrows indicate a visible seam where the tissue layers were brought together, but seams are no longer visible by 10 hours later. hpa = hours post-amputation. Scale bars: 50µm (B), 100 µm (C), 25 µm (E).

We first amputated juvenile worms transversely below the pharynx, so that only the outer epidermis (but not the pharyngeal epithelium) would be impacted by the injury. Consistent with previous reports (cite), each fragment showed a rounded wound edge within 24 hours, suggesting successful closure. Head fragments formed a pointed tail within 3 days (red arrowhead) and regained their original shape within 5 days. The tail fragments took longer to regenerate but formed a visible blastema within 5 days (white arrowhead) and regained their original form within 10 days (Figure 2B). This temporal trajectory reflects the regenerative challenge each fragment faces; the tail fragments need to replace more cell types than their head counterparts. Although the brightfield imaging suggested that wound sites had closed within 24 hours, we wanted to assess the cellular dynamics of the re-epithelialization process.

To track the dynamics of the outer epidermis during wound closure, we stained fragments with the cytoskeletal dye SiR-actin at intermediate time points during the first 10 hours post-amputation (hpa). Wounds from both head and tail fragments largely closed within 10 hpa, and both showed a strong anisotropic enrichment of actin at the wound periphery, suggesting the formation of a supracellular structure (Figure 2C-D). These observations appear consistent with a process of radial constriction, where actin polarizes towards the wound edge and cells collectively pull towards the center of the wound to minimize the size of the gap. Some wound edges displayed a smooth, continuous actin cable (yellow arrowheads), implying the formation of an actomyosin purse-string^31,36^, but others appear puckered (red arrowheads), suggesting more of a general constrictive mechanism. Both head and tail fragments closed their wounds at a similar rate; head wounds were more open than tail wounds at 3 hpa, but by 6 hpa we found no difference between the two conditions (Figure 2D). Despite closure of the wound sites by 10 hpa, we observed visible differences in wound edges between 10 and 20 hpa. Closed wounds at 10 hpa had visible seams where the epidermis was brought together (yellow arrows), but by 20 hpa there was no detectable seam or wound edge (Figure 2E). This suggests a sustained re-epithelialization process even after initial gap closure, where the epidermal cells continue to remodel their junctions and re-establish a seamless epithelium.

To assess whether minor wounds utilize the same mechanisms of actomyosin contractility during re-epithelialization as the transversely amputated wounds above, we performed ‘pinch’ wounds, where fine-tipped forceps were used to pinch through the outer epidermis (Supplemental Figure 1A). These wounds closed faster than wounds at transverse amputations and seemed to use a different cellular mechanism. Instead of contracting across the wound gap, we observed the formation of polarized protrusions within 5 minutes post-amputation, with the wound edges making contact and closing up within 1 hour (Supplemental Figure 1A). This is likely due to the much smaller distance between wounded epidermal edges; in a pinch wound, the edges remain adjacent to each other, facilitating quick re-epithelialization, but a transverse amputation causes a comparatively large wound gap that requires more time to close.

Together, these results show that wounds that disrupt only one epithelial layer use shared mechanisms of closure seen in other systems across contexts of regeneration, development, and repair. Minor wounds with small gaps can close quickly and without the formation of supracellular structures, whereas large amputations require the constriction of epithelial cells across the wound gap. We next looked at the dynamics of closure in more complex wounds, specifically injuring both the outer epidermis and the pharyngeal epithelium.

### Heterotypic epithelial bridges mediate the first step in multi-epithelial wound closure

To investigate mechanisms of wound closure involving multiple epithelial layers, we performed injuries that damage both the outer epidermis and the pharynx. We first cut animals in the sagittal plane, separating the left half from the right, and observed wounds over a 10-day time course of regeneration (Figure 3A-B). Interestingly, these animals were able to mostly restore their original shape within 5 days, rounding themselves up without the formation of a visible blastema (Figure 3B). We therefore asked how the epithelial layers facilitate the morphallactic processes of wound closure. Using SiR-actin labeling, we amputated worms in the sagittal plane and observed the formation of striking, long cytoplasmic protrusions that extend across the wound edge in the first 10 hpa. These protrusions originated from both the outer epidermis and the pharyngeal epithelium, connecting across the middle of the wound (Figure 3C). To confirm that these cells were epithelial in nature, we utilized the *EpiA:RFP* transgenic line^52^ and detected protrusions in labeled cells within both the outer epidermis and the pharyngeal epithelium (Supplementary Figure 2A-B). We were unable to visualize the basal side of cells forming these bridges, but the extensions protruded from the apical surface of the cells - suggesting that these were new structures and not movement of the homeostatic basal protrusions.

**Figure 3.**
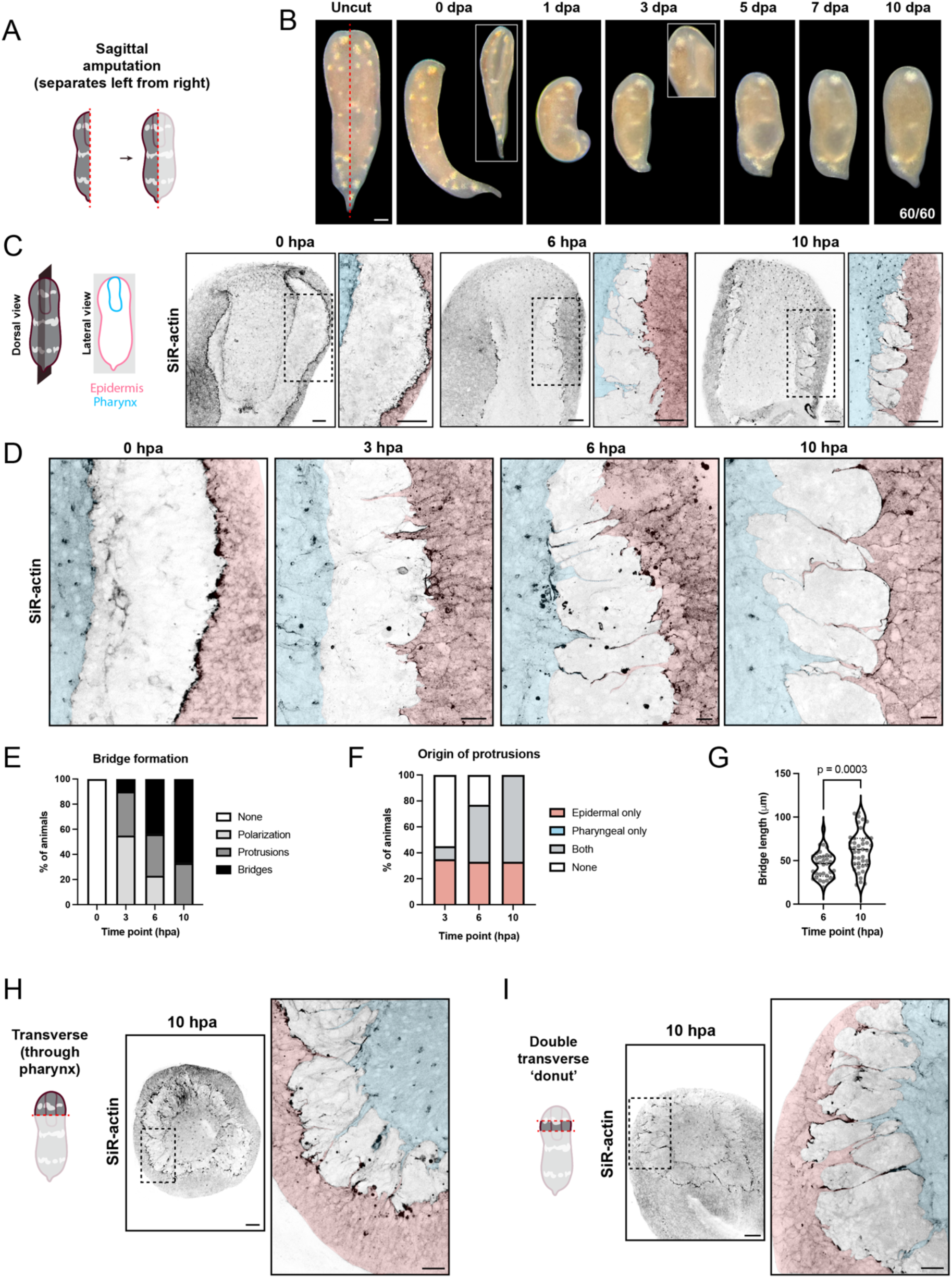
Injuries that damage both epithelial tissues form long, heterotypic protrusions connecting the epithelial layers across the wound edge. (A-B) Schematic (A) and representative brightfield time course (B) showing regeneration of *H. miamia* following sagittal amputation. (C) Representative images of wound closure following sagittal amputation between 0-10 hours post-amputation. (D) High-magnification images showing polarization, protrusion, and formation of heterotypic bridges during the first 10 hours of wound closure. (E-G) Quantifications of heterotypic bridge formation (E), origin of protrusions (F), and combined bridge length (G) over the first 10 hpa. Data represent 8 worms per condition and time point (E-F) and 30-40 bridges per time point (G); comparison by unpaired *t-*test. (H-I) Representative images demonstrating the presence of heterotypic bridges in single (H) and double (I) transverse amputations through the pharynx. In all high-magnification images, the outer epidermis is pseudocolored in pink and the pharyngeal epithelium is pseudocolored in blue for clarity. Scale bars: 50µm (B-C, H-I), 10 µm (C, zoomed-in box), 5 µm (D), 20 µm (H-I, zoomed-in box).

In sagittal wounds, epidermal cells showed signs of actin polarization within the first 3 hours post-amputation (hpa), with small filopodial protrusions (5-10 µm) appearing in the outer epidermis oriented towards the wound edge (Figure 3B). Long, thin extensions (>15 µm) emerged around 6 hpa, forming more robustly by 10 hpa and connecting to projections originating from the pharyngeal epithelium (Figure 3D-E). This temporal pattern was consistent across animals, and 67% of fragments displayed at least one connected bridge within 10 hpa (Figure 3E). Notably, protrusions were always detected first in the outer epidermis, and protrusions from the pharyngeal epithelium were only visible after their counterparts formed in the epidermis (Figure 3F). Corresponding with the number of bridges increasing between 6 and 10 hpa, the average bridge length increased significantly between time points (Figure 3G). This stepwise process (polarization, extension, connection) is consistent with a model of collective cell migration^54,55^, but the lengths of these bridges (∼30-100 µm) are reminiscent of specialized projections such as cytonemes or tunneling nanotubes (which have been reported to reach up to 200 µm *in vivo*^56–58^). To the best of our knowledge, structures like these heterotypic bridges have not been reported in wound closure literature, and we therefore sought to further investigate this phenomenon.

The difference between transverse, contraction-based closure and sagittal, bridge-based closure could be attributed to a variety of differences between the amputations other than the dual epithelial layers, including (but not limited to): angle or orientation of amputation, size of the wound, or the extent of regeneration required downstream of closure. We next asked whether heterotypic bridges occurred in transverse amputations further up the anterior-posterior axis of the animal, upon injuries that impact both epithelial layers by cutting through the pharynx. These wounds require closure in the same plane as the transverse injuries reported above, but now include damage to the pharyngeal epithelium. When we performed single and double transverse amputations through the pharynx, we observed robust heterotypic bridges between the epidermis and the pharyngeal epithelium, similar to those seen in sagittal amputations (Figure 3H-I). This bridge-based mechanism of closure is therefore not specific to the size or the irregular wound geometry that comes from a sagittal amputation, but appear to instead be a more general mechanism by which to close wounds when both epithelial layers are damaged.

Although these heterotypic bridges are unique in presentation, we reasoned that there must be a substrate below these protrusions acting as a physical track for their extension. The extracellular matrix (ECM) has been shown to be a key regulator of wound healing, acting as a dynamic substrate on which cells can crawl and extend to complete re-epithelialization^42,59,60^. Given the known role of collagen IV in wound closure^43,60–62^, we identified a putative ortholog in the *H. miamia* transcriptome and made a custom antibody (see Methods). Using this antibody, we performed immunofluorescence staining to assess the localization of collagen IV and detected its expression solely under the pharyngeal epithelium (Supplemental Figure 2C-D). Next, we asked whether collagen IV was remodeled during wound closure as the pharyngeal epithelium is extending its protrusions. We performed immunofluorescence on 10 hpa sagittal fragments and observed strong co-localization of collagen IV specifically underneath pharyngeal extensions. Collagen signal was never detected in areas without protrusions, suggesting that nascent collagen is being deposited underneath the pharyngeal epithelium as it is forming wound-induced bridges (Supplemental Figure 2E). The same pattern was observed after double transverse amputation, suggesting a shared mechanism regardless of wound geometry (Supplemental Figure 2F). These data implicate collagen IV as a substrate for epithelial bridges, likely providing physical support for protrusions as they extend across the wound site. Although our antibody specifically labels the collagen underneath the pharyngeal epithelium, we speculate that a similar process is occurring underneath protrusions coming from outer epidermal cells.

Taken together, these findings reveal the presence of heterotypic cellular bridges between epithelial layers during wound closure, which emerge within the first 10 hours post-amputation. Tissue-specific collagen is remodeled during the extension of pharyngeal protrusions, and bridges are observed across different amputation geometries that injure multiple epithelial layers. We therefore hypothesized that these heterotypic interactions act as an intermediate step in large-scale multi-epithelial wound closure, possibly minimizing tissue loss and/or maintaining tissue identity prior to full re-epithelialization. To the best of our knowledge, this phenomenon appears to be a novel healing mechanism, and we sought to understand how these heterotypic interactions relate to the rest of the wound healing process.

### Wound closure involving multiple epithelial tissues utilizes distinct but temporally overlapping cellular mechanisms

The bridges described above represent a transient phase in wound closure, emerging within the first 10 hours post-amputation (hpa). In order for the animal to heal fully, the sagittal fragments must undergo continued remodeling to re-establish their original shape. Our regeneration time course demonstrated that sagittal amputations in *H. miamia* do not form a blastema, but instead slowly round up over the course of 70 hours (Figure 3B). To gain further insight into the cellular processes driving this morphogenetic movement, we stained sagittal fragments with SiR-actin over the course of 70 hours post-amputation (hpa) (Figure 4A). We observed that these animals healed through a two-step process: as described above, the outer epidermis first connects to the pharyngeal epithelium within the first 10 hpa, followed by the posterior end closing itself up through what appears to be a combination of epithelial crawling and zippering. Both of these cellular movements are well-described in other systems, and here we see a similar gradual closure of the outer epidermis as it meets and re-epithelializes with the epidermis from the other side of the animal.

**Figure 4.**
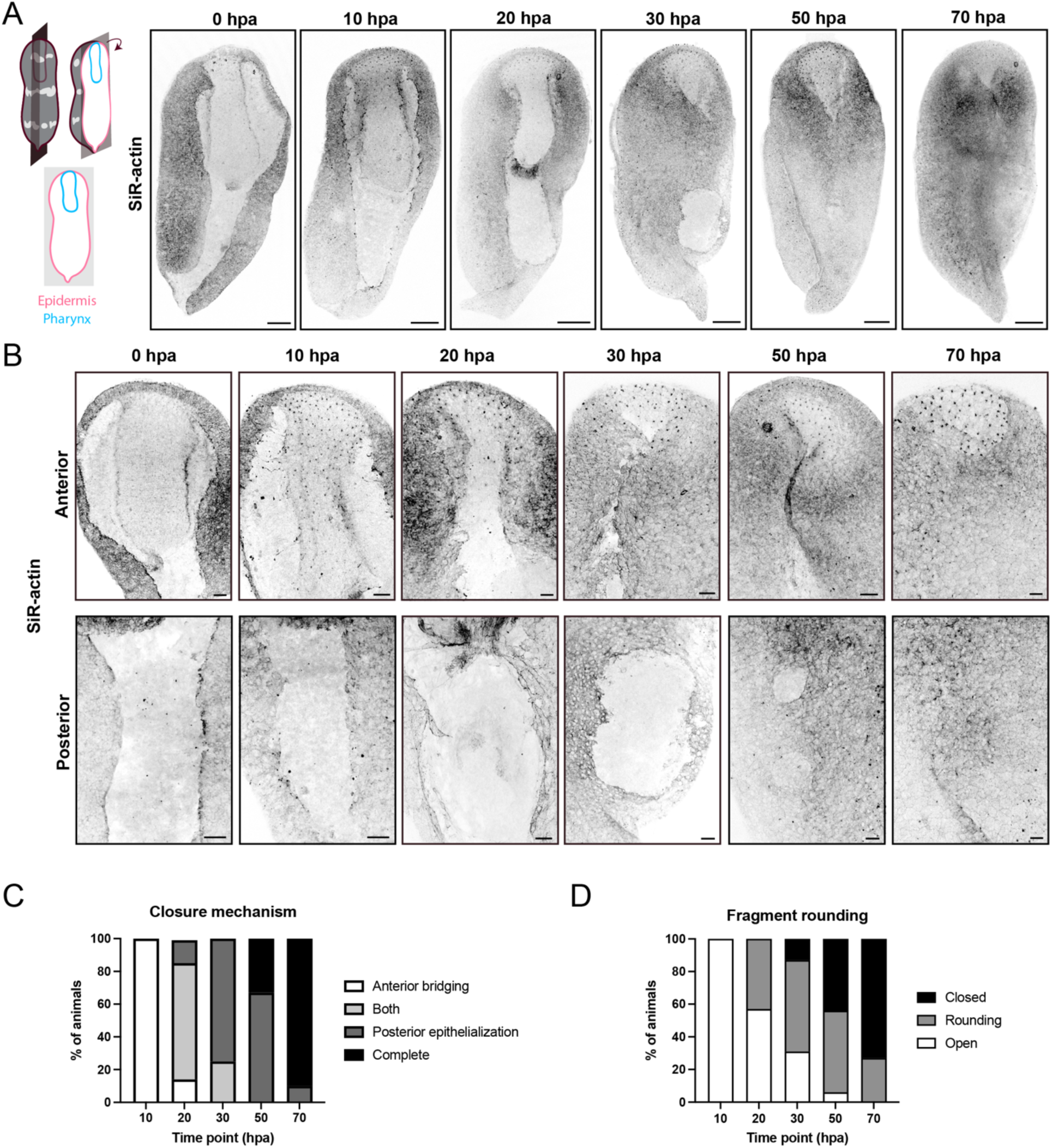
Sagittal wounds heal through a two-step process, combining epithelialization with whole-body rounding to restore the fragment’s original morphology. (A) Schematic and time course showing epithelial tissue dynamics in sagittal wound closure during the first 70 hours post-amputation (hpa), using SiR-actin labeling. (B) Higher-magnification images demonstrating the overlapping mechanisms of epithelialization in the anterior and posterior parts of the fragment. Anterior tissues form heterotypic protrusions described in Figure 3, and the posterior wound edges gradually close through homotypic re-epithelialization. (C-D) Quantifications of closure mechanisms (C) and fragment rounding (D). Data represent n=12 animals per time point. hpa = hours post-amputation. Scale bars: 100 µm (A), 20 µm (B).

As the posterior ends are closing, sagittally-cut animals round their bodies up to re-form their cylindrical shapes. The two processes we observexd (heterotypic bridges and homotypic re-epithelialization) occur in a stepwise but overlapping sequence, where all fragments first form visible bridges, then display both bridges and posterior closure, and finally end with just posterior re-epithelialization (Figure 4B-C). All animals had an open body shape between 0-10 hpa, characterized by a splayed pharynx and separated dorsal and ventral halves. At 20 hpa, we began to see rounding of the fragments, with the outer epidermal edges curling over the pharynx towards each other. While some animals closed as early as 30 hpa, virtually all animals completed this closure by 70 hpa (Figure 4B-D). Here, closure is defined by the original dorsal and ventral edges completely meeting at the seam of the wound, forming a cylinder with the pharynx on the inside and the epidermis on the outside. We speculate that this rounding process is aiding the posterior re-epithelialization by bringing the wound edges together.

To rule out that this dual closure mechanism was unique to the sagittal amputation, we next asked whether similar processes occurred in transverse injuries that form heterotypic bridges. We amputated animals both once (single transverse) and twice (donut) through the pharynx and tracked regeneration. The single transverse fragments followed a similar regeneration trajectory as their counterparts amputated below the pharynx (Figure 2B), although the differences in fragment size correlated with different temporal dynamics (Supplemental Figure 3A). The donut fragments formed blastemas on both anterior and posterior ends, regenerating all missing structures despite the severity of the injury (Supplemental Figure 3B). To track the dynamics of wound closure in these amputations, we fixed animals at intermediate time points and stained fragments with SiR-actin. Unlike the sagittal fragments, the transverse fragments didn’t zipper after the formation of bridges, but instead contracted their outer epidermis over the wound site in a process that appeared similar to the single-epithelial transverse amputations (Supplemental Figure 3C-D, Figure 2B).

These observations suggest that while the exact mechanism of wound closure is determined by a combination of factors, injuries involving both epithelial layers follow a coordinated two-step process. First, heterotypic protrusions bridge the wound gap between the two epithelial layers, followed by the outer epidermis re-epithelializing with itself. We next asked how cell and tissue contractility was involved in these mechanisms of wound closure, seeking to combine the characterization of these processes with functional perturbations.

### Muscle contraction is necessary for both heterotypic wound closure and downstream morphogenetic movements

Contraction of muscle at the wound site is an essential component of wound closure in both vertebrates and invertebrates, driving physical and biochemical events through the generation of mechanical force^22,26,29,63^. Because of the close proximity of epithelial cells to underlying muscle in *H. miamia* (Figure 1G), we next asked whether muscle contraction was involved in wound closure. *H. miamia* have an extensive network of muscle throughout their bodies (Figure 5A), and we utilized Tropomyosin immunofluorescence to visualize how muscle was reorganized during single and dual epithelial wound closure. In transverse wounds (both head and tail), muscle was largely absent from the wound site itself but was instead present at the periphery of the wound edge as it shrank (Figure 5B). In contrast, sagittal wounds contain parenchymal muscle fibers that link the outer epidermis to the pharyngeal epithelium, coming together as the animal rounds up (Figure 5C). We hypothesized that this difference in muscle organization might influence the closure mechanisms used by each fragment, and asked whether muscle contraction was required for each type of wound closure.

**Figure 5.**
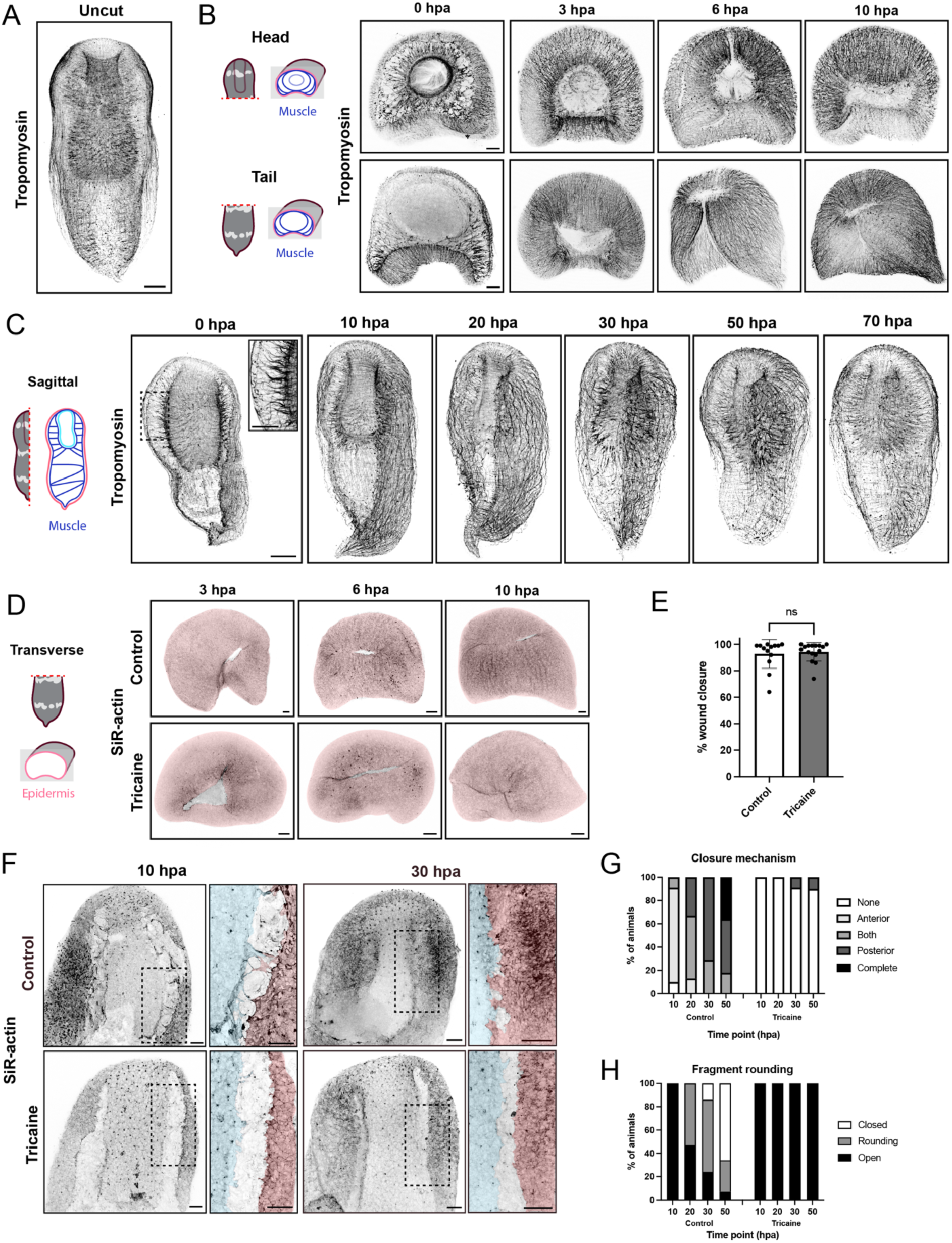
Muscle contraction is dispensable for traditional transverse amputations but required for the formation of heterotypic bridges during sagittal wound closure. (A-C) Images showing the organization of muscle in uncut worms (A), transverse wound closure (B), and sagittal wound closure (C), visualized by immunofluorescence staining for tropomyosin. (D-E) Representative images (D) and quantification (E) of wound closure in control and tricaine-treated animals after transverse amputation, labeled with SiR-actin. (F) Representative images of wound closure in control and tricaine-treated animals after sagittal amputation, labeled with SiR-actin. In D and F, the outer epidermis is pseudocolored in pink and the pharyngeal epithelium is pseudocolored in blue for clarity. (G-H) Quantification of closure mechanism (G) and fragment rounding (H) in control and tricaine-treated animals after sagittal amputation. Data represent n=12 animals per condition, per time point; quantification in (E) by unpaired *t-*test. ns=not significant, hpa = hours post amputation. Scale bars: 100 µm (A,C), 50 µm (B,D,F).

To test the functional role of muscle contraction, we anesthetized amputated animals in 0.1% tricaine, which blocks neural action potentials^64^ and effectively inhibits muscle contraction in *H. miamia* (Video 1). In the presence of tricaine, single epithelial wound closure in transverse amputations occurs at the same rate as controls (Figure 5C-D), indicating that the epidermis is capable of constricting and re-epithelializing without the aid of muscle contraction. Animals with multi-epithelial wounds, on the other hand, are incapable of forming heterotypic bridges without muscle contraction (Figure 5E-G). We observed some polarization and formation of small protrusions from the epidermis and pharyngeal epithelium, but no bridges ever formed in tricaine-treated animals (Figure 5F). The requirement of muscle contraction for the formation of heterotypic bridges was consistent across all multi-epithelial injuries (Supplemental Figure 4A-B), suggesting a shared mechanism.

We performed time courses with tricaine treatment to determine whether muscle contraction was only necessary for the initiation of the heterotypic bridges or whether muscle contraction was needed for both their formation and maintenance (Supplemental Figure 4C). Animals treated with tricaine for 10 hours, then placed in sea water for 20 hours, closed wounds at a similar rate to controls. In contrast, animals that were in sea water for 10 hours and then placed in tricaine for 20 hours were indistinguishable from animals treated with tricaine for all 30 hours (Supplemental Figure 4D-E). This indicates both that persistent muscle contraction is necessary for the maintenance of heterotypic bridges, and that animals can fully recover after anesthetization.

Interestingly, muscle contraction is only required for the participation of the outer epidermis in forming heterotypic bridges. In the presence of tricaine, the pharyngeal epithelium still formed extensions and spread to try and cover the wound site (Supplemental Figure 4F-G). While the exact function is not yet clear, these data implicate muscle contraction as a necessary driver of multi-epithelial wound closure. This contraction may be involved in generating tension, inducing mechanical or biochemical signals, and/or physically bringing the wound edges together to allow contact between the epithelia. Without muscle-generated mechanical force, the outer epidermis cannot generate or maintain long protrusions to form heterotypic bridges, and the wound closure process stalls and collapses.

Together, these findings highlight the importance of muscle contractility in mediating cellular events during multi-epithelial wound closure. To distinguish between the role of tissue-scale mechanical forces generated by muscle and the cell-intrinsic forces required for protrusive activity and wound closure, we next assessed the role of actomyosin contractility in wound closure.

### Types of wound closure are differentially regulated by actomyosin contractility

Actomyosin contractility is generated through the interactions of non-muscle myosin with actin, producing mechanical forces within cells that enable cytoskeletal rearrangements and cellular movements^65–67^.

While actomyosin contractility is broadly implicated in wound closure across species, we wanted to know how its role varied across injury types in *H. miamia*. We treated amputated animals with the inhibitor blebbistatin, which inhibits the ATPase activity of myosin and blocks actomyosin contractility^68,69^. In the presence of blebbistatin, transverse wounds (which only disrupt the outer epidermis) closed at comparable speeds to control conditions but appeared to switch towards more crawling-dominant mechanisms (Figure 6A-C): instead of the contractile rings seen in control fragments, we detected protrusions around the wound edge and saw individual cells crawling across the wound site (Figure 6C). These data indicate that protrusion-based mechanisms are sufficient to accomplish wound closure in blebbistatin-treated conditions.

**Figure 6.**
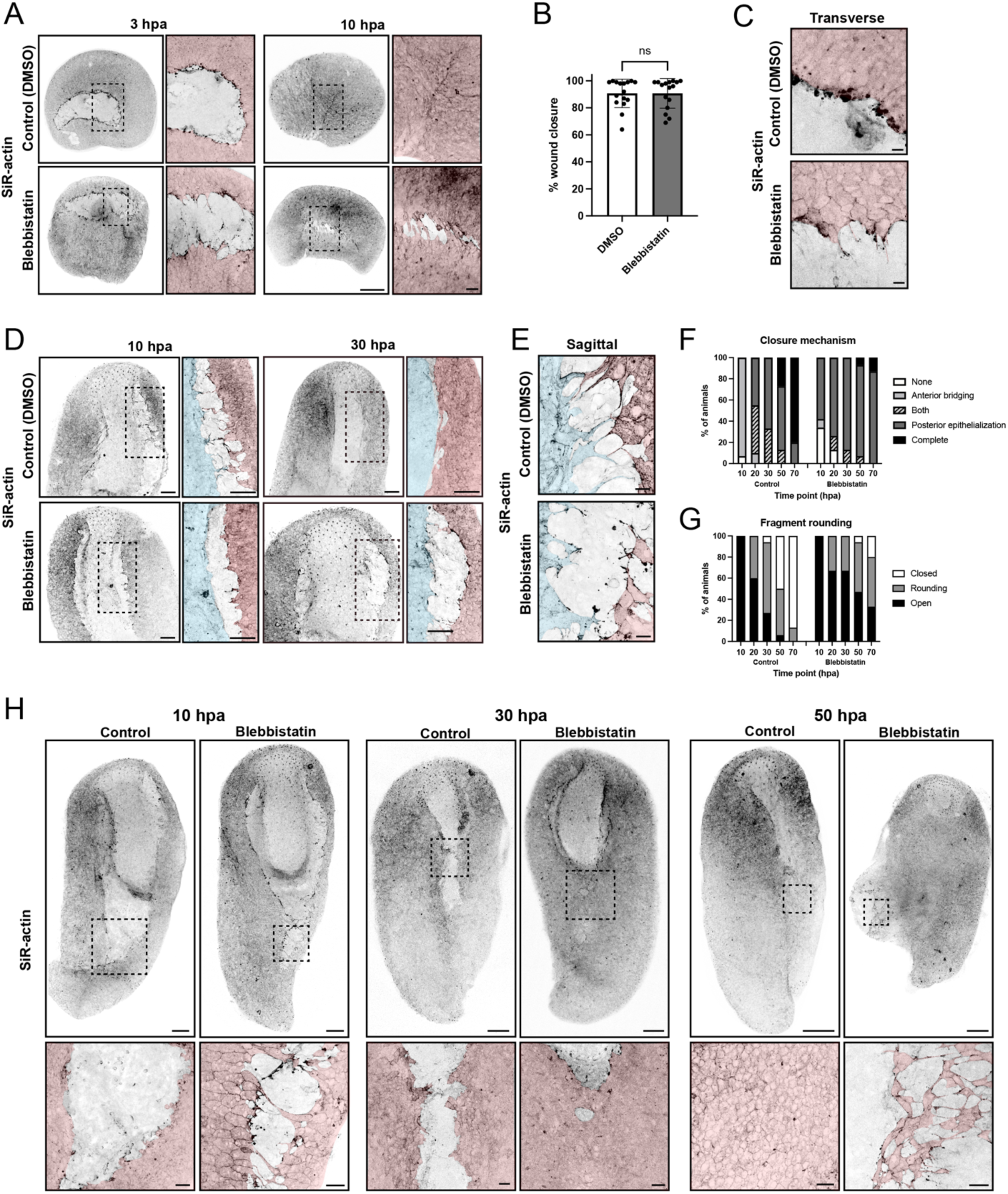
Actomyosin contractility is differentially utilized across wound closure mechanisms. (A-B) Representative images (A) and quantification (B) of wound closure in control and blebbistatin-treated animals after transverse amputation, labeled with SiR-actin. (C) High-magnification images showing the difference between control and blebbistatin-treated wound edges in transverse amputations. (D) Representative images of wound closure in control and blebbistatin-treated animals after sagittal amputation. (E) High-magnification images showing heterotypic bridges in control animals and the lack thereof in blebbistatin-treated animals. (F-G) Quantification of closure mechanism (F) and fragment rounding (G) in control and blebbistatin-treated animals after sagittal amputation. (H) In high-magnification images, the outer epidermis is pseudocolored in pink and the pharyngeal epithelium is pseudocolored in blue for clarity. Data represent n=12-16 animals per condition, per time point; comparison in (B) by unpaired *t-*test. ns=not significant, hpa = hours post amputation. Scale bars: 50 µm (A,D), 10 µm (C,E), 100 µm (H), 20 µm (H, zoomed-in box).

We next asked how actomyosin contractility was involved in multi-epithelial wound closure. In contrast to the mild phenotype in single epithelial wounds, we observed significant closure defects in the blebbistatin-treated animals. Blebbistatin treatment eliminated the formation of stable heterotypic bridges in sagittal fragments at 10 hpa (Figure 6D). We observed short protrusions emerging from both the outer epidermis and pharyngeal epithelium, but their orientation appeared more random than control fragments, forming multiple protrusions rather than connected bridges (Figure 6E, Supplemental Figure 5A). These data indicate that actomyosin contractility is essential for the extension of long protrusions across the wound edge to form heterotypic bridges. This contractility may be important in the stabilization of protrusions as they extend, the application of pulling forces to bring the epithelia together, and/or the stabilization of nascent adhesions at heterotypic interfaces^34,39,66,70–72^. Without this contractility, small protrusions can form, but they ultimately lack the necessary force to form bridges.

Contrasting with the inhibition of heterotypic bridges, we observed an acceleration of epidermal re-epithelialization in blebbistatin-treated animals. In control conditions, the zippering up of the posterior end began around ∼20-30 hours after amputation, after the formation of heterotypic bridges at the anterior end (Figure 4D). In contrast, animals treated with blebbistatin showed posterior closure as early as 10 hpa, even in the absence of heterotypic bridges (Figure 6F-H). This phenotype is consistent with that seen in other systems, in which the reduction of junctional tension ‘loosens’ the epithelial sheet and allows epidermal cells to move more freely relative to each other^30,68,73^. By 20 hpa, blebbistatin-treated animals showed robust closure of the outer epidermis, but no heterotypic bridges and no rounding of the pharyngeal epithelium. Prolonged treatment with blebbistatin for 50 hours, however, led to reduced tissue integrity in the epidermis, manifesting first as holes between cells and leading to eventual epidermal ruptures (Figure 6H). The presumed decrease in junctional tension that enabled accelerated cell spreading also prevented the maintenance of strong junctions between cells, resulting in a weakened epidermal layer that was unable to withstand the mechanical demands of healing and regeneration.

Lastly, we assessed whether this phenotype was reversible and if the epidermis could re-establish its integrity after accelerated re-epithelialization. We placed blebbistatin-treated animals in fresh sea water after a 30-hour incubation for either 20 (50 hpa time point) or 40 (70 hpa time point) hours, comparing directly to control animals and animals left in blebbistatin for the full time course (Supplementary Figure 5B). Animals that were placed in fresh ASW for 20h were rounder than their counterparts that remained in blebbistatin, suggesting a re-entry into the typical morphogenetic trajectory (Supplementary Figure 5C-D). Animals that were placed in fresh ASW for 40 hours were able to completely recover, resembling the control animals at 70 hpa (Supplementary Figure 5C-D). This suggests that while prolonged blebbistatin treatment will lead to a loss of epidermal integrity, the animals will restore junctional tension after removal of the drug, continuing the re-wound closure process.

## DISCUSSION

In this study, we demonstrate that the acoel *Hofstenia miamia* uses context-dependent wound healing, employing complementary and overlapping mechanisms of re-epithelialization to repair wounds prior to whole-body regeneration. Wounds disrupting only one tissue use rapid and conserved closure mechanisms, similar to wound closure in other species. In control conditions, these wounds primarily rely on radial constriction of the epidermis, but in contractility-inhibited conditions the epidermal cells shift mechanisms and close through crawling. Multi-epithelial wounds close through a combination of novel heterotypic bridges and homotypic epithelial zippering, requiring both actomyosin-generated tension and muscle contraction to drive coordinated tissue movements. In conditions of reduced global contractility, other closure mechanisms are transiently upregulated, but ultimately fail to restore long-term tissue integrity unless contractility is restored. Mechanisms of wound closure in *H. miamia* are therefore not uniform or discrete processes, but are instead highly dynamic and cooperative, shifting in response to both intrinsic and extrinsic factors.

The heterotypic bridges described here are, to the best of our knowledge, a novel wound-induced response. Heterotypic contacts have been described in other systems as mechanisms by which tissue boundaries are maintained during development^75–79^, but filopodial matching that has been observed^79^ appears distinct from these bridges both in size and duration. Instead of fast filopodial probing between proximal cells to form the proper adhesive complexes, the bridges observed in *H. miamia* are long (>50 um) and visible for at least 10 hours. At present, we cannot rule out the possibility that these bridges are dynamic due to a lack of live imaging, but their length paired with the clear progression of formation (polarization of actin, followed by short protrusions, then connected bridges) suggests some temporal stability. Although they differ in presentation, it’s possible that these heterotypic contacts are functioning similarly to maintain tissue borders. During regeneration, injuries that damage both epithelial layers must preserve both tissue identity and orientation, ultimately restoring the ‘tube within a tube’ anatomy. Because we do not yet have tissue-specific labeling to differentiate between the epithelial layers, we don’t know how the bridges resolve during secondary closure. However, the framework presented here will set the stage for future studies, including better visualization of cellular interactions, identification of adhesion and migration machinery, the role of signaling pathways, and more.

When assessing the dynamics of these heterotypic interactions, we observed that the outer epidermis requires muscle contraction to form stable protrusions, but the pharyngeal epithelium can spread across the wound even in the absence of epidermal contributions. This phenotype could be due to intrinsic differences in the epithelia; for example, perhaps the pharyngeal epithelium has intrinsic cell motility machinery that is primed for migration regardless of external forces. In the animal’s undamaged configuration, the pharyngeal epithelium is surrounded by rings of pharyngeal muscle. It’s possible that when that muscle is damaged, the epithelial cells are now able to spread and crawl across the wound site. In contrast, the outer epidermal cells may be more stable and require assistance from muscle contraction to bring the two layers close enough to form stable adhesions. The differences between the epithelial responses could also be due to their extrinsic interactions. The epithelial cells in *H. miamia* are anatomically unique, lacking a flat basal surface as is typically observed in most animal epithelia and instead forming long projections that end in close proximity to muscle. Electron microscopy in acoels has failed to identify a visible basement membrane underneath the outer epidermis^74^, but the composition of extracellular matrix (ECM) in *H. miamia* has not been studied. It is therefore unclear whether the ECM is adhering to protrusions in a punctate fashion, if there is ECM surrounding muscle fibers, if there is a thin layer of ECM in between cellular protrusions, or any other organization. Using a custom antibody in this study, we demonstrate that there is tissue-specific collagen underneath the pharyngeal epithelial cells, and that collagen is present underneath their wound-induced protrusions. We did not observe this collagen underneath the outer epidermis, but a similar mechanism could be occurring with a different extracellular matrix protein, perhaps one whose remodeling is mechanosensitive. Differences in ECM stiffness and composition could affect the mechanical response of each tissue, as well as differences in regular tensile loads. It has also been shown in other systems that wound-induced muscle contraction triggers the release of matrix remodeling proteins^42,59^, which could be necessary for epidermal cells to extend far enough to reach their pharyngeal epithelial counterparts. Future studies will need to further characterize the *H. miamia* ECM and its dynamics to distinguish between these hypotheses.

The difference in wound closure strategies across injury types raises several questions about the connection between healing and regeneration. It seems probable that different forms of wound closure correlate with different cellular and molecular environments in regenerating fragments, whether through biochemical and/or mechanical events associated with re-epithelialization. Future work will be required to test whether different closure strategies have functional consequences for downstream tissue regeneration, whether healing and regeneration are distinct and nonoverlapping biological processes, and whether other species capable of whole-body regeneration utilize distinct mechanisms to heal different types of wounds.

## MATERIALS AND METHODS

### *H. miamia* strains and husbandry

Adult worms were housed in artificial sea water (ASW) (37 ppt, pH 7.70-8.00) in plastic boxes within 21°C incubators. Embryos were collected from adult boxes and placed into petri dishes for development. Once hatched, juvenile worms were transferred to tanks for post-embryonic growth. All animals were fed twice a week; adult worms were fed brine shrimp (Artemia, Brine Shrimp Direct) and juvenile hatchlings were fed rotifers (*Brachionus plicatilis*, Reed Mariculture). *EpiA:RFP* worms were generated as previously described^52^.

### Amputations and sample preparation

Juvenile worms were isolated and starved for 1-2 weeks prior to use to minimize autofluorescence of ingested food. For regeneration experiments, animals were placed in 1mg/mL ethyl 3-aminobenzoate, methanesulfonic acid (“tricaine”, stock solution 5mg/mL of ASW) (Thermo Fisher, AC118000100) in ASW in a petri dish for 3-5 minutes until they stopped swimming, then amputated using a scalpel. After amputation, fragments were transferred to a dish containing fresh ASW until their designated time point. For fixation, animals were placed in 4% paraformaldehyde (PFA) (Thomas Scientific, C993M31) in ASW for 1 hour at room temperature. The PFA was removed and animals were washed with PBST (PBS + 0.1% Triton-X-100) three times prior to immunofluorescence staining. For pinch wounds, animals were grasped with fine-tip forceps, then allowed to recover in ASW prior to fixation. For transverse and donut cuts, animals were cut once or twice transversely, then transferred to fresh ASW until their designated time point. For single transverse wounds, animals were cut transversely once again, right above the previous cut, to remove the wound site for imaging. These fragments were then immediately fixed in 4% PFA in ASW.

### Small molecule inhibitors and treatment with anesthetics

For small molecule inhibitor experiments, animals were first amputated as described above, then flushed with ASW to remove tricaine prior to incubation in designated inhibitors. Animals were left in inhibitor for 3-70 hours, depending on the experiment. Blebbistatin (Sigma-Aldrich, B0560) was dissolved in DMSO per manufacturer instructions, then diluted in ASW for use at 25µM. DMSO control treatments used an equal volume of DMSO as Blebbistatin treatments. For tricaine time point experiments, animals were left in 0.5mg/mL tricaine in ASW for 3-50 hours.

### Immunostaining

For antibody staining, all steps were carried out using an orbital shaker. Fixed animals were washed in PBST for 3×5 minutes, then placed in blocking solution (PBST + 10% goat serum (Thermo Fisher, 16210072)) for one hour at room temperature. Animals were then placed in primary antibody in block solution for two nights at 4°C. Primary antibody was washed out with PBST, 6×20 minutes. Animals were then blocked once again for one hour at room temperature, then incubated with secondary antibody (1:800) (Thermo Fisher, A-11011) in blocking solution overnight at 4°C. The next day, secondary antibody was washed out using PBST, 6×20 minutes. Finally, animals were incubated in SiR-actin (1:1000) in PBST (Cytoskeleton, CY-SC001) for 90 minutes at room temperature prior to mounting. All samples were mounted on double cover slips using Prolong Glass Antifade Mountant (Thermo Fisher, P36980). If animals were only stained with SiR-actin, they were placed in SiR-actin in PBST for 90 minutes at room temperature, then mounted immediately. The custom *H. miamia* Tropomyosin antibody^80^ (Rabbit IgG) was used at 1µg/mL Where relevant, Hoechst (1:500) (Invitrogen, H3570) was used to counter-stain nuclei.

### Generation of custom collagen IV antibody

Rabbit polyclonal antibodies were custom-made by GenScript through their PolyExpress immunization protocol, using a peptide sequence from *H. miamia* Collagen IV (CPPGAKGNDGEPGEKG). This peptide was expressed in *E. coli* using the provided sequence and then used to immunize two rabbits. Two antibodies were generated, purified, and validated in-house by Western Blot with the protein immunogen.

### Microscopy and image processing

All images were collected using 10x/0.25x (dry), 20x/0.75x (dry), or 63x/1.40 (oil) fluorescent objectives on a Leica DMI SP8 inverted confocal microscope, equipped with Leica Application Suite software. Images were processed using Image J/FIJI software.

### Quantifications

#### Length of epithelial basal protrusions

To quantify the average length of basal protrusions in outer epidermal and pharyngeal epithelial cells, images were taken of labeled cells in *EpiA:RFP* animals. The Measure function in FIJI was used to quantify the length of each protrusion, tracing from the end of the protrusion to where it meets the body of the cell. For each cell type (pharyngeal epithelium vs. outer epidermis), *n*=15 cells from n=5 animals (total = 45 cells) was measured.

#### Percentage of closure in transverse amputations

For quantification of wound closure in transverse cuts, images were taken of both the wound edge and the open edge of each sample, using FIJI to measure the open area of each side. The percentage of wound closure was determined by dividing the open area of the wound edge by the total open area, then subtracting that from 1. For each time point and fragment (head vs. tail), measurements were collected from *n*=8 animals.

#### Percentage of closure in sagittal amputations

To quantify the percentage of wound closure after sagittal amputations, images of SiR-actin-stained fragments were imported into FIJI. Because of mounting imperfections, only one side of each animal was measured, choosing the side with the clearest signal for the most accurate quantification. 6-12 animals were quantified per time point and condition. Within FIJI, the Measure function was used to quantify: 1) the length of the pharynx and 2) the cumulative distance of connected tissue between the epidermis and the pharyngeal epithelium. The percentage of wound closure was determined by dividing the closed distance by the total pharynx length. For each time point, measurements were collected from *n*=12 animals.

#### Bridge length

To quantify the length of heterotypic bridges, images of SiR-actin-stained fragments were taken at each time point, then imported into FIJI for analysis. Using the Measure function, bridges were quantified by drawing a line along the membrane extensions protruding between cell types. Cell bodies were not included in this measurement. Because of the technical challenge of robustly identifying where one protrusion met the other, the combined bridge length was reported instead of individual cell contributions. For each time point, measurements were collected from *n*=30-40 animals.

#### Morphological descriptions

To report the morphological characteristics of healing fragments, images of SiR-actin-stained fragments were taken at each time point. For bridge formation, animals were binned into four categories: none (no visible polarization of the epithelial layers and no protrusions), polarization (visible polarization of cell membranes towards the wound edge), protrusions (at least one membrane extension <15um in length), or bridges (at least one connected heterotypic bridge). For protrusion origin, animals were assessed for visible protrusions coming from only the outer epidermis, only the pharyngeal epithelium, both, or neither. Rounding was assessed by looking at the shape of the animal, binning animals into either open (no visible rounding), rounding (partway through the rounding process), or closed (where the outer edges of the fragments have met, re-forming a cylindrical body plan). The values plotted for each morphological trait were the representative percentage of total fragments, ranging from *n*=10-45 animals (exact values are reported in accompanying figure legends).

### Graphing and statistical analyses

All graphs were generated using GraphPad PRISM. Data with two or more factors were analyzed using two-way ANOVA with Dunnett’s multiple comparisons test. Data comparing changes in one condition over different time points were analyzed by one-way ANOVA with Dunnett’s multiple comparisons test. A two-tailed unpaired *t*-test with Welch’s correction was used to compare data between two independent groups. All statistical tests are stated in figure legends. *P* values were obtained using GraphPad PRISM and used to determine the level of varying significance between experimental groups; groups were considered different when *P*<0.05.

## ACKNOWLEDGEMENTS

APK is a Fellow of the Jane Coffin Childs Memorial Fund for Medical Research. This project was supported by grants from the National Institute of General Medical Sciences of the NIH to MS: R35GM128817 and R35GM153252. Thanks to members of the Srivastava Lab for feedback on the manuscript.

## AUTHOR CONTRIBUTIONS

A.P.K conceived the project, performed experiments, quantified and analyzed data, and wrote the manuscript under the supervision of M.S. Both authors discussed the results and edited the manuscript.

## SUPPLEMENTARY FIGURES AND TABLES

**Supplementary Figure 1.**
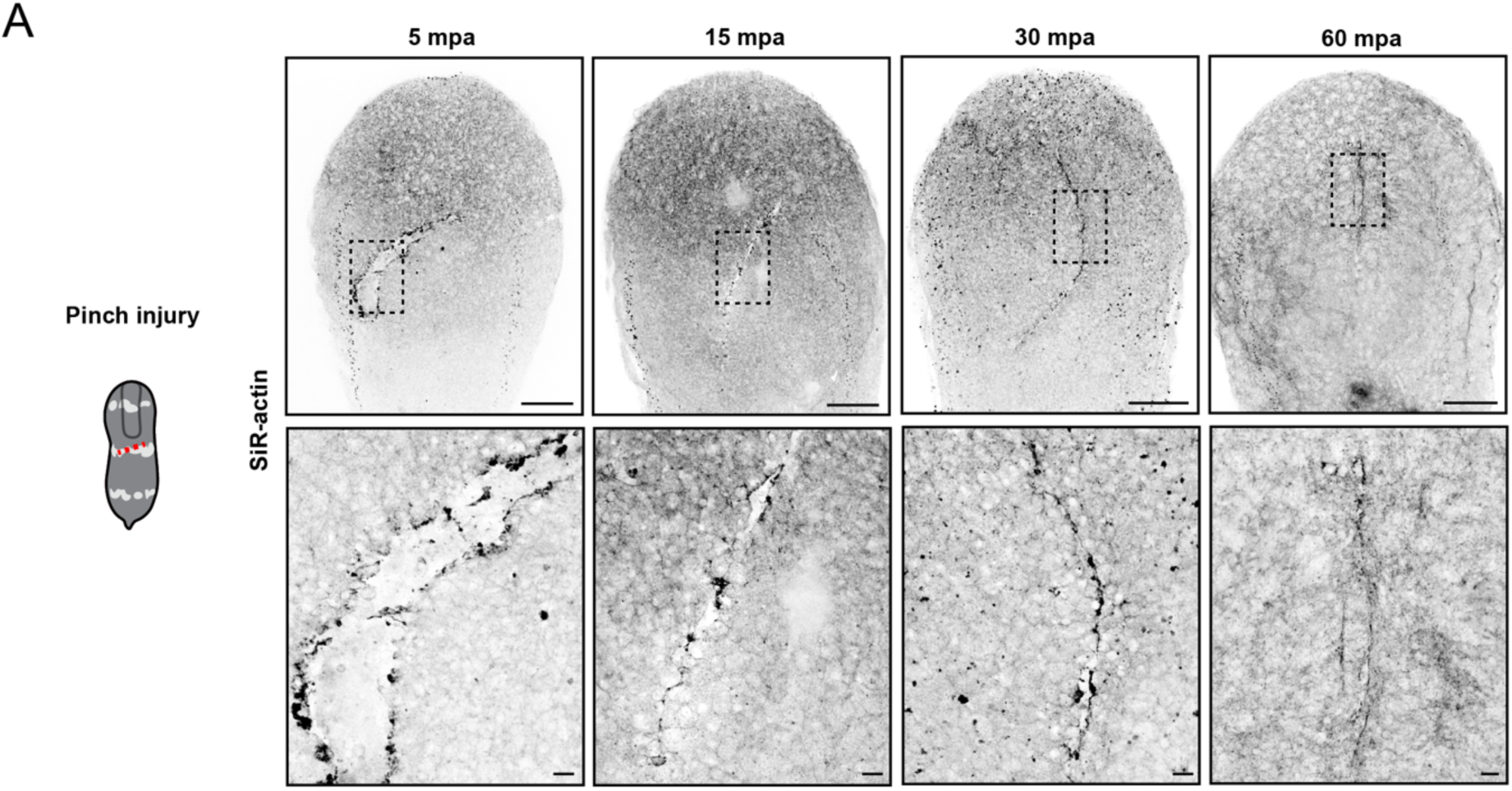
Pinch wounds heal rapidly, utilizing cell protrusions to bring the wound edges together. (A) Schematic and representative images showing wound closure following an epidermal pinch injury, labeled with SiR-actin. Scale bars: 100 µm (A, top row), 10 µm (A, zoomed-in box). mpa = minutes post amputation.

**Supplementary Figure 2.**
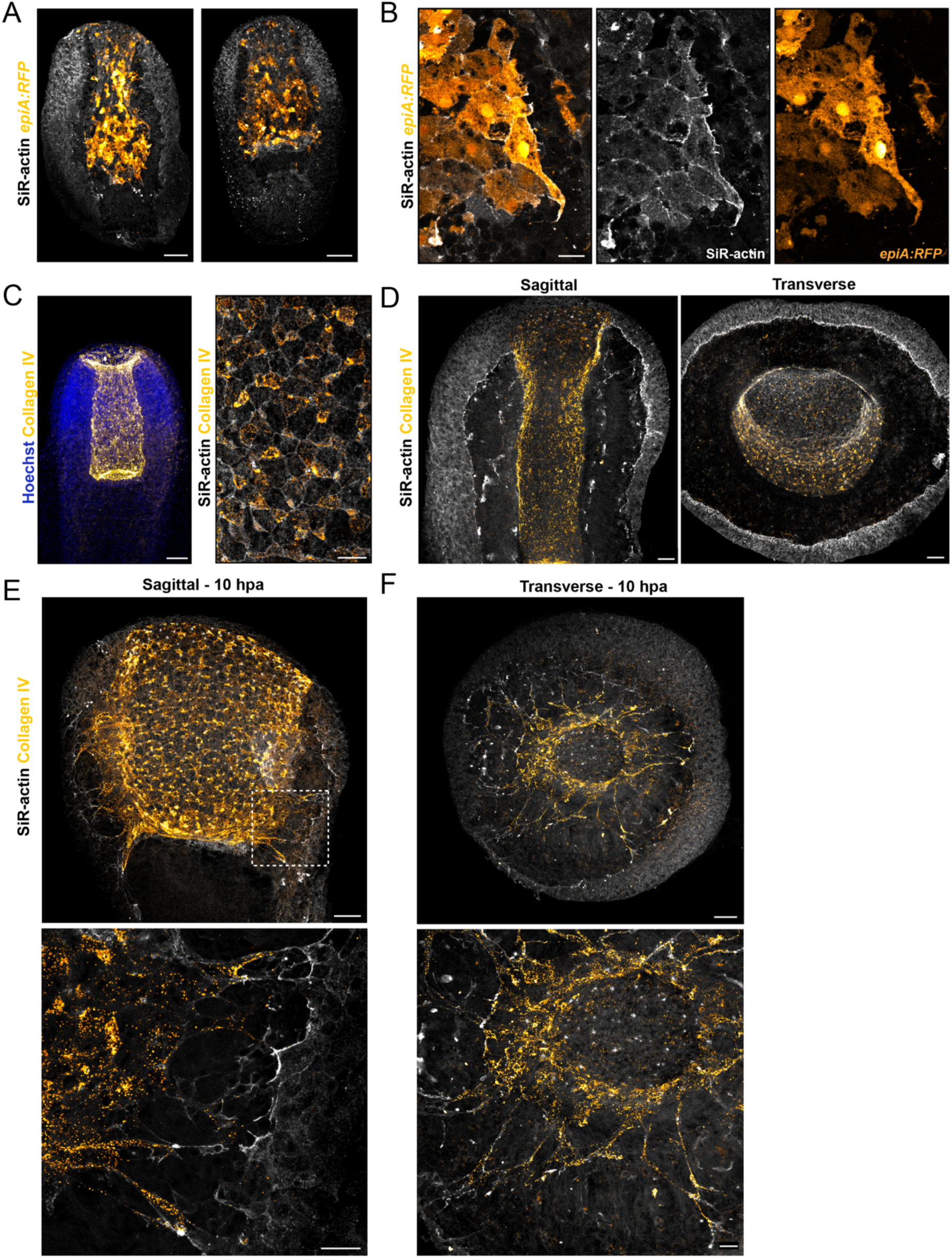
Collagen IV is visible underneath pharyngeal, but not epidermal, protrusions during wound closure. (A-B) *EpiA:RFP* transgenic animals cut in the sagittal plane have mosaic expression of RFP in both the epidermis and the pharyngeal epithelium (A), and these cells are part of the SiR-actin-labeled population that is forming protrusions during wound closure (B). (C-F) Immunofluorescence staining using a custom Collagen IV antibody (see Methods) showing localization underneath the pharyngeal epithelium in uncut (C) and 10 hours post-amputation (hpa) (D-F) fragments. Scale bars: 50µm (A, C (left), D, E-F (top)), 10µm (B), 20µm (C (right), E-F (bottom)).

**Supplementary Figure 3.**
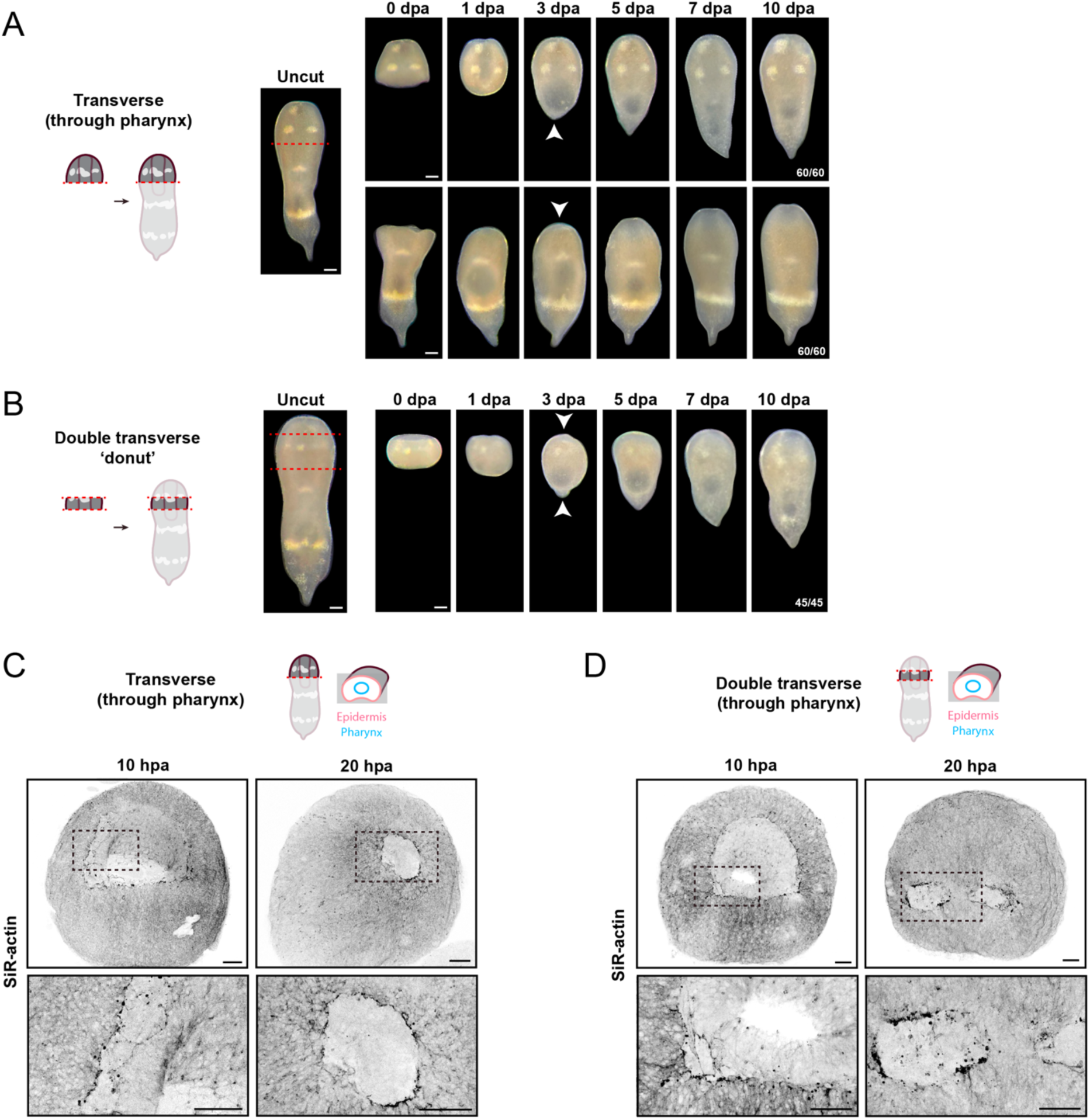
Transverse amputations that injure the pharyngeal epithelium heal through a similar two-step mechanism. (A-B) Schematic and representative brightfield time course showing regeneration of *H. miamia* following single (A) and double (B) transverse amputation. (C-D) Representative images showing wound closure following single (C) and double (D) transverse amputation, labeled with SiR-actin. Scale bars: 100µm (A-B), 50µm (C-D).

**Supplementary Figure 4.**
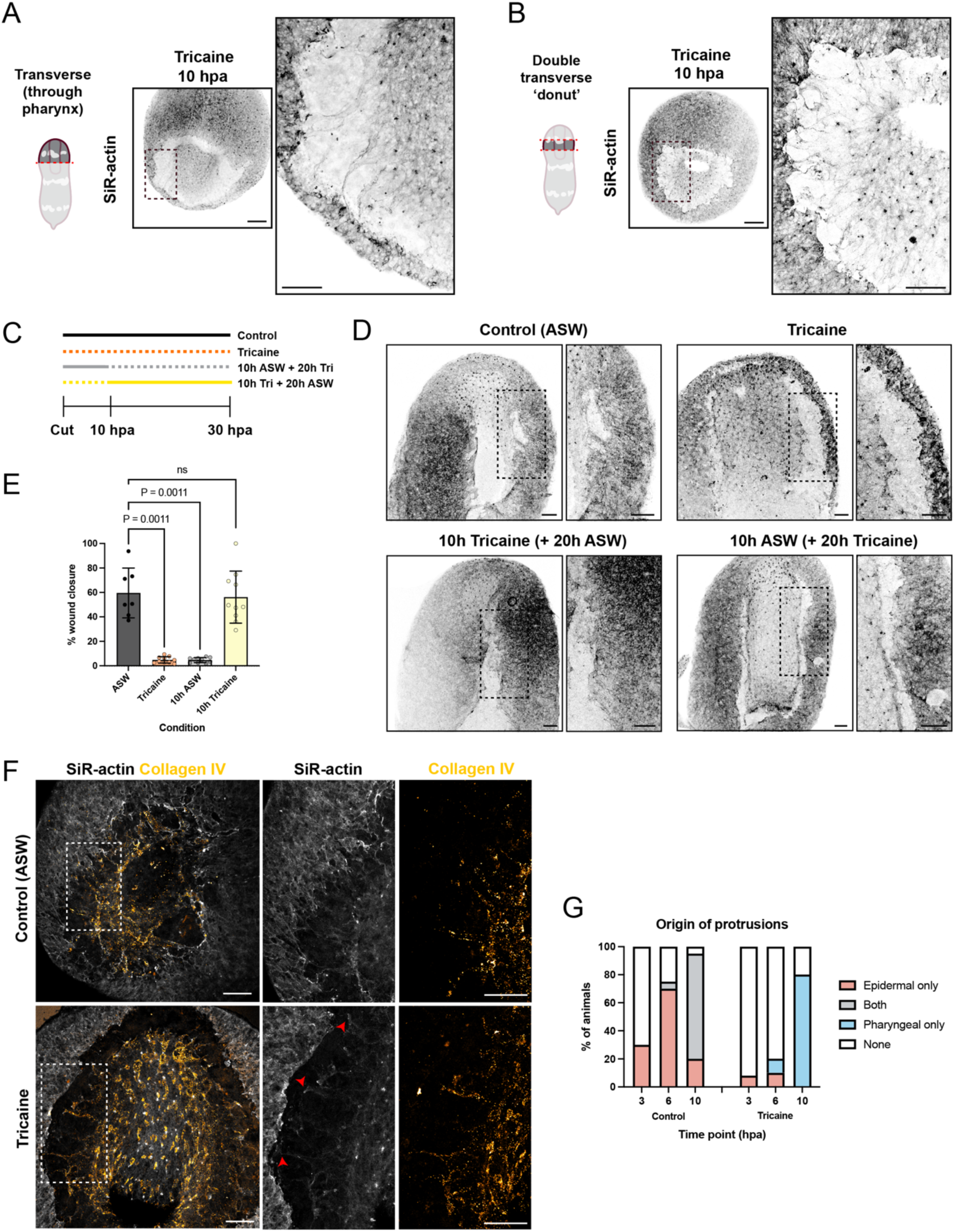
Muscle contraction is required for epidermal protrusions during heterotypic bridge formation. (A-B) Representative images showing pharyngeal protrusions after single (A) or double (B) transverse amputation in tricaine-treated conditions. (C) Schematic of tricaine timing experiment. (D) Representative images showing wound closure in control, tricaine, 10h tricaine (+ 20h ASW), or 10h ASW (+ 20h tricaine) conditions. (E) Quantification of wound closure across tricaine conditions. Data represent n=8 worms per condition and show mean ± s.e.m. Comparisons by one-way ANOVA with Dunnett’s multiple comparisons test; ns=not significant. (F) Representative images showing Collagen IV localization underneath pharyngeal extensions following transverse amputation in control and tricaine-treated conditions. (G) Quantification of protrusion origin in control and tricaine-treated animals. Data represent n=7-10 animals per condition, per time point. Scale bars: 100µm (A,B), 50µm (A,B (zoomed-in boxes), D,F).

**Supplementary Figure 5.**
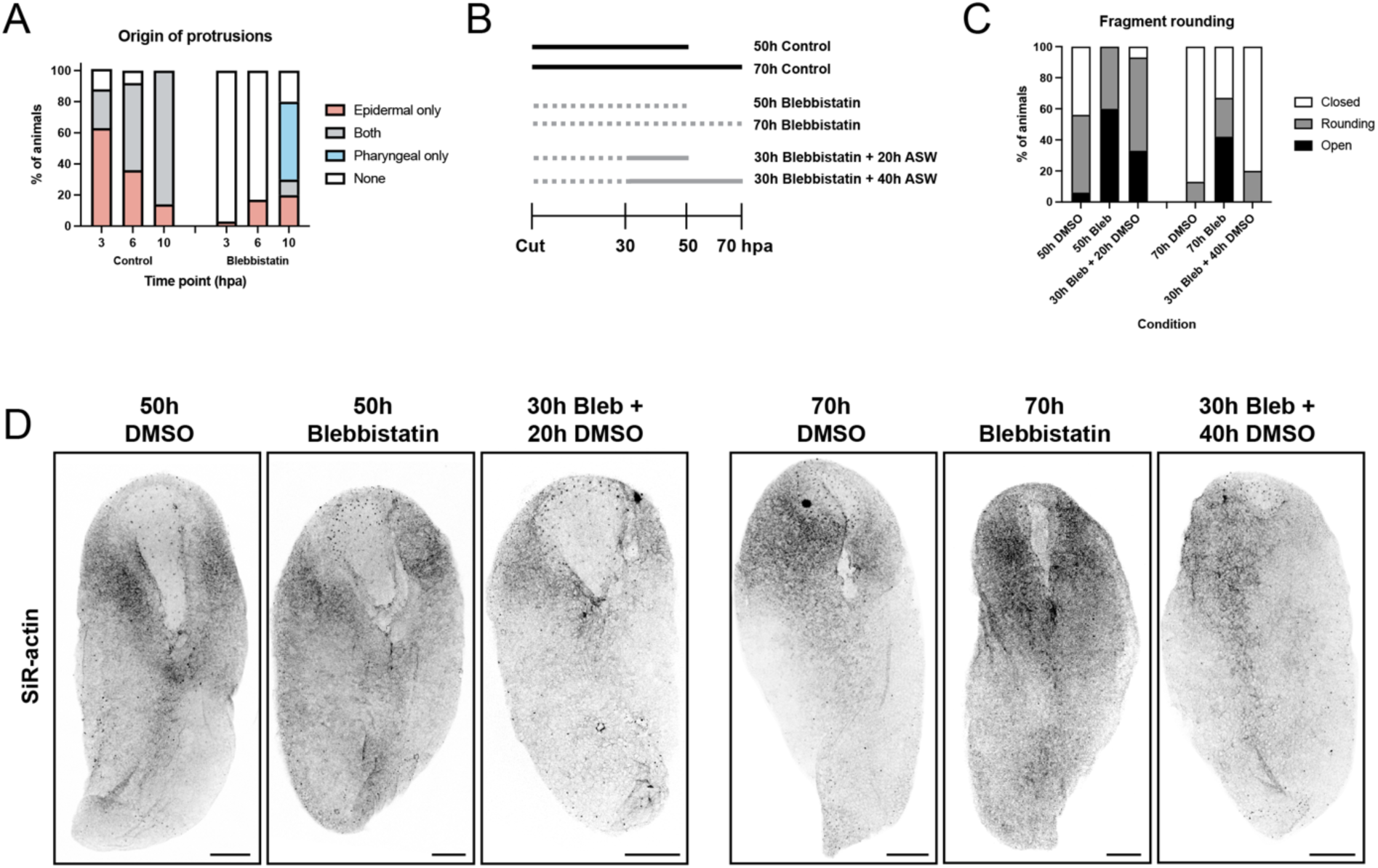
Blebbistatin treatment is reversible. (A) Quantification of protrusion origin in control and blebbistatin-treated animals. (B) Schematic of blebbistatin wash-out experiment. (C-D) Quantification (C) and representative images (D) of fragment rounding following control and blebbistatin treatments. Data represent n=8 animals per condition, per time point. Scale bars: 100µm (D).

**Supplementary Video 1. Tricaine slows muscle contraction in *H. miamia*.**

Animals are anesthetized in 1mg/mL tricaine in ASW.

